# Neuregulin-1 Attenuates Myocardial Ischemia/Reperfusion Injury by Activating the UCP2/PINK1/LC3B-mediated Mitophagy

**DOI:** 10.1101/2024.10.08.617324

**Authors:** Xin-Tao Li, Xin-Yue Li, Tian Tian, Wen-He Yang, Yi Cheng, Kai Su, Xi-Hua Lu, Mu Jin, Fu-Shan Xue

## Abstract

**BACKGROUND:** Ischemia/reperfusion (I/R) injury may significantly affect the treatment outcomes and prognosis of patients with acute myocardial infarction following coronary artery recanalization. Available evidence suggests that neuregulin-1 (NRG-1) can provide a protection against myocardial I/R injury and is involved in various cardioprotective interventions by potential regulation of mitophagy. However, the molecular mechanisms linking NRG-1 and mitophagy remain to be clarified. This experiment aimed to determine whether NRG-1 postconditioning attenuated myocardial I/R injury through the regulation of mitophagy and to explore the underlying mechanisms.

**METHOD:** Both an *in vivo* myocardial I/R injury model of rats and an *in vitro* hypoxia/reoxygenation (H/R) model of H9C2 cardiomyocytes were applied. NRG-1 treatment was conducted immediately after I/R or H/R intervention. In the *in vivo* experiment, cardioprotective effects of NRG-1 were determined by infarct size, cardiac enzyme and histopathologic examinations. The potential downstream pathways and molecular targets of NRG-1 were screened by the RNA sequencing and the Protein-Protein Interaction Networks (PPI). The expression levels of mitochondrial uncoupling protein 2 (UCP2) and mitophagy-related protein in both the I/R myocardium and H/R cardiomyocytes were measured by immunofluorescence staining and Western blots. The activation of mitophagy was observed with the transmission electron microscopy (TEM) and JC-1 staining.

**RESULTS:** The KEGG and GSEA analyses showed that the mitophagy-related pathways were enriched in the I/R myocardium treated with NRG-1, and UCP2 exhibited a significant correlation between mitophagy and interaction with PINK1. Meanwhile, the treatment with mitophagy inhibitor Mdivi-1 significant eliminated the cardioprotective effects of NRG-1 postconditioning *in vivo*, and the challenge with UCP2 inhibitor genipin could also attenuate the activating effect of NRG-1 postconditioning on mitophagy. Consistently, the *in vitro* experiment using H9C2 cardiomyocytes showd that NRG-1 treatment significantly up-regulated the expression levels of UCP2 and mitophagy-related proteins, and activated the mitophagy, whereas the challenge with small interfering RNA (siRNA)-mediated UCP2 knockdown abolished the effects of NRG-1.

**CONCLUSIONS:** NRG-1 postconditioning can produce a protection against the myocardial I/R injury by activating mitophagy through the UCP2/PINK1/LC3B signaling pathway.

## Introduction

Acute myocardial infarction (AMI) is one main reason of global population deaths. It is generally believed that restoring blood perfusion to ischemic myocardium is one of the most effective treatments for AMI^1^. However, this restoration of blood flow may cause ischemia/reperfusion (I/R) injury, which may account for up to 50% of the final myocardial infarct area and significantly worsen the treatment outcomes and prognosis of patients^2,3^. Therefore, both investigating the potential mechanisms of myocardia I/R injury and exploring effective strategies to mitigate the I/R injury are essential for improving the outcomes of patients with AMI.

Neuregulin-1 (NRG-1), belonging to the epidermal growth factor family, serves as an intercellular signaling molecule. It is primarily localized in coronary blood vessels and endothelial cells. Available evidence indicates that NRG-1 plays a crucial role in regulating myocardial remodeling, myocardial regeneration, and cardiac electrical signaling^4–6^. It has been shown that the hypoxia/reoxygenation (H/R) pretreatment for the endothelial cells can increase the NRG-1 level in the culture medium, which produces a significant protection against the H/R injury of cardiomyocytes^7,8^. Most important, recent work reveals that NRG-1 postconditioning can replicate the cardioprotective effects of ischemic postconditioning by activating the RISK pathway^9^. These findings suggest that NRG-1 plays an important role in the cardioprotective benefits of various interventions. Nonetheless, the precise mechanisms involved still require further clarification.

Because mitochondria serve as the primary energy source of cardiomyocytes, maintaining the normal structure and function of mitochondria is essential for the physiological activity of cardiomyocytes. The normal homeostasis and function of mitochondria are highly dependent on the mitochondrial quality control (MQC), which involves the prompt removal of damaged mitochondria and the reutilization of mitochondrial components. As a critical stage in MQC, mitophagy is a selective mitochondrial degradation process which is particularly crucial for maintaining mitochondrial function and homeostasis^10^. The increasing evidence indicates that the regulation of mitophagy plays an important role in the protection of myocardial injury with different interventions^11–13^. Especially, the cardioprotection of ischemia preconditioning is negated by the knockout of the mouse PINK1 gene, which is a crucial molecule involved in the process of mitophagy^14^. Furthermore, NRG-1 treatment can protect against the H/R cardiomyocytes by decreasing mitochondrial membrane potential (MMP)^15^, and the reduction in the MMP is a crucial factor triggering the PINK1-mediated mitophagy. However, it remains unclear whether regulating mitophagy is one of potential mechanisms for cardioprotection of NRG-1. Consequently, this experiment aimed: (1) to determine whether NRG-1 postconditioning could provide a protection against myocardial I/R injury; and (2) to explore whether PINK1-induced mitophagy was involved in the cardioprotective effects of NRG-1 postconditioning and its underlying molecular mechanism.

## Methods

### Data Availability

The supporting data for the results of this research can be obtained from the corresponding author upon a reasonable request.

### Animals

All the experiment protocols were reviewed and approved by the Animal Care and Use Committee of Beijing Friendship Hospital (No. 22-1011, October 11, 2022). Male Sprague–Dawley rats (weighted 220-270 g, aged 5-7 weeks) were supplied by the Beijing Vital River Laboratory Animal Technology Co., Ltd. (Beijing, China). They were housed in a specific pathogen-free laboratory at the Animal Experimental Center of Beijing Friendship Hospital under the standard conditions with controlled temperature (24±2 °C), humidity (40-60%) and a 12-h light/dark cycle. The experiment procedures were conducted according to the Guidelines for Animal Experimentation of our institute and Animal Research—Reporting of *In Vivo* Experiments (ARRIVE) guidelines.

### Establishment of *in vivo* rat model with myocardial I/R injury

The *in vivo* myocardial I/R injury model was established, according to the methods previously described^16^. Following anesthetized with an intraperitoneal injection of 2% tribromoethanol 10 mg/kg (Sigma, USA), the animal received the tracheotomy and tracheal cannula, and then were ventilated using a small animal ventilator with respiratory rate of 55 breaths/min and tidal volume of 2.5 ml/100 g body weight. The electrocardiogram and hemodynamic variables were continuously recorded by the electrodes placed subcutaneously on the limbs and the right common carotid artery cannula with a animal monitor (Techman Instrument Co., Ltd., Chengdu city, China). By a left thoracotomy in the fourth intercostal space, the left anterior descending (LAD) coronary artery was ligated with a slipknot to induce the regional myocardial ischemia. Following a 30-minute ischemia, the ligature was released for 120 minutes of reperfusion. Ischemia was confirmed by the pale and cyanotic myocardial tissue along with an obvious ST segment elevation on electrocardiogram^17^.

### Animals grouping and treatments

Using a computer-generated random number table, animals were assigned into five groups to receive the different interventions: (1) Sham-operated (Sham) group, the animals received the shame surgical procedures without a ligation of the slipknot around the LAD artery for 150 min; (2) ischemia/reperfusion injury (IRI) group, the animals received the I/R surgical procedures with a 30-min ischemia and a 120-min reperfusion; (3) NRG-1 postconditioning (NRG-1) group, the animals received the I/R surgical procedures and the treatment of NRG-1 3 μg/kg (MedChemExpress, Monmouth Junction, USA), which was given through the internal jugular vein immediately following the end of ischemia^18^; (4) Mdivi-1 treatment (Mdi-1) group, a mitophagy inhibitor known as Mdivi-1 (MedChemExpress) was administered via an intraperitoneal injection with a dosage of 10 mg/kg immediately before the ligation of LAD artery^19^, and then the animals were underwent the same interventions as the NRG-1 group; (5) Genipin treatment (Gen) group, genipin (MedChemExpress), an inhibitor of mitochondrial uncoupling protein 2 (UCP2), was intraperitoneally given at a dose of 30 mg/kg biweekly for two times before the I/R surgical procedures^20^, and then the animals were underwent the same interventions as the NRG-1 group.

### Cells and treatments

The rat H9C2 cardiomyocytes for the *in vitro* experiment were cultured and propagated as described previously^21^. Then, the H/R intervention was carried out as described in previous work^22^. Briefly, cardiomyocytes were exposed to hypoxic conditions with 1% O_2_ and 37 °C for 6 h and then were reoxygenated at 37 °C for 12 h in a normoxic incubator. The NRG-1 treatment on cardiomyocytes was performed as previously described^15^. That is, the NRG-1 (MedChemExpress) was added to the culture medium at a concentration of 200 ng/mL immediately following the hypoxic intervention, after which reoxygenation was conducted.

### Cells Transfection

The small interfering RNA (siRNA) against the UCP2 and negative control siRNA were purchased from the OBiO Technology (Shanghai, China). The H9C2 cardiomyocytes were transfected with the complex containing the UCP2 siRNA (50 nM) and Lipofectamine^TM^ RNAiMAX transfection reagent (Thermo Fish Scientific, Waltham, Massachusetts) for 48 h in the light of the manufacturer’s protocols. The siRNA targeting sequences used in this study as follow:

Forward:5’-AGAGCACUGUCGAAGCCUACATT-3’,

Reverse:5’-UGUAGGCUUCGACAGUGCUCUTT-3’.

### Evans blue and TTC staining

The double-staining of Evans blue and 2, 3, 5-triphenyltetrazolium chloride was applied to determine the myocardial infarct size as previously described^23^. In summary, following the reperfusion for 120 min, the LAD artery was re-ligated and 2% Evans blue dye (Sigma-Aldrich, USA) was injected through the right common carotid artery to demarcate the ischemic area at risk (AAR) and normally perfused region of heart. Following the animal was euthanized by increasing the depth of anesthesia and intravenous administration of 10% potassium chloride at a dose of 100 mg/kg, the heart was excised and rapidly frozen at −80 °C condition for 15 min. The heart located beneath the ligation site was sliced along its long axis into 1 mm sections. Next, these heart pieces were incubated in 1% TTC solution at 37 °C condition for 15 min to determine the infarcted myocardium area. An investigator who was blind to the grouping assignment analyzed and quantified the infarct size of heart slices by the ImageJ software (infarcted myocardium area/AAR×100%).

### Serum creatine kinase isoenzyme (CK-MB) and cardiac troponin I (cTnI) assessment

Following a 120-min reperfusion, 3-ml blood sample was collected through the right common carotid artery in each rat. The serum supernatant was extracted and then serum CK-MB and cTnI levels were measured using the ELISA kits (Elabscience Biotechnology Co., Ltd, Wuhan, China) in adherence to the manufacturer’s instructions.

### Myocardial histopathology

Following the conclusion of experiment, the hearts were removed and the I/R myocardial samples were immersed immediately in 4% paraformaldehyde. Subsequently, the samples were embedded in paraffin and sliced into the 5-μm sections. After staining the slides with hematoxylin and eosin (HE), the pathological changes of the I/R myocardium were observed and captured using a light microscope.

### RNA sequencing

At the end of experiment, the rats from the IRI group and NRG-1 group were euthanized by increasing the depth of anesthesia followed by an intravenous injection of 10% potassium chloride 100 mg/kg. The hearts were removed and the I/R myocardial samples were immersed in RNA*later*™ Stabilization Solution (Invitrogen, Carlsbad, USA) overnight at 4 °C. The total RNA of the I/R myocardium was isolated and purified using the TRIzol reagent (Invitrogen, Carlsbad, USA) following the instructions of manufacturer.

The amount and quality of RNA in each sample were quantified using NanoDrop ND-1000 (NanoDrop, Wilmington, USA). The RNA integrity was determined by the Bioanalyzer 2100 (Agilent, Santa Clara, USA) and confirmed by the electrophoresis with denaturing agarose gel. Both the RNA-sequencing (RNA-seq) and the subsequent analysis were conducted by the Lianchuan Biotechnology Co., Ltd (Hangzhou, China).

### Transmission Electron Microscopy (TEM)

As previously described^11^, both the mitochondria and mitophagy in the I/R myocardium observe by the TEM. Briefly, at the conclusion of experiment, the hearts were excised and rinsed with the ice-cold PBS. Then the I/R myocardium was cut into the blocks of 1 mm^3^ and sealed in the electron microscopy fixation solution (Servicebio, Wuhan, China) under the light proof condition with temperature of 4 °C. Subsequently, the tissue samples were embedded with acetone and embedding agent mixture, and then sectioned into ultrathin slices and stained using uranyl acetate and lead citrate. The tissue sections were examined and photographed using a transmission electron microscope (HITACHI, Tokyo, Japan).

### Immunofluorescence staining

As previously described^24^, the I/R myocardial samples were embedded with paraffin to cut into the multiple 5-μm sections, and the antigen retrieval process was applied by heating sections in the EDTA (pH 8.0) to 95 °C for 15 min. Following a 2-h incubation with 5% BSA for blocking, the tissue slides were incubated at 4 °C condition overnight with the primary antibody of anti-LC3B (1:200, ABclonal Technology) or anti-UCP2 (1:50, Santa Cruz Biotechnology, Santa Cruz, USA). The Alexa 488 secondary antibody (1:400, Abcam, United Kingdom) was incubated at ambient temperature for 1 h. As for the cells’ staining, H9C2 cardiomyocytes were seed in 12-well plates on slides for subsequent experiment. At the end of experiment, the slides were fixed in 4% paraformaldehyde at room temperature for 20 min and incubated with 0.5% TritonX-100 for 5 min. After blocking with 10% FBS and 0.1% bovine serum albumin in the PBS at room temperature for 30 min, the slides were incubated with anti-LC3B primary antibody (1:50, ABclonal Technology) and MitoTracker^TM^ dyes (Invitrogen, USA). The secondary antibody labeling was same as before. The images were captured using an IX-83 confocal microscope (Olympus, Tokyo, Japan).

### Assessment of mitochondrial membrane potential

The assessment of mitochondrial membrane potential (MMP, ΔΨm) was conducted utilizing the JC-1 Mitochondrial Membrane Potential Assay Kit from MedChemExpress. When the ΔΨm elevates, JC-1 aggregates develop and emit red fluorescence (Ex/Em=585/590 nm). Conversely, when the ΔΨm reduces, JC-1 exists as monomers and emits green fluorescence (Ex/Em=510/527 nm). An increase in the formation of JC-1 aggregates signifies mitochondrial membrane depolarization, leading to a decrease in ΔΨm, which in turn initiates mitophagy^25^. H9C2 cardiomyocytes were cultured in 12-well plates containing slides and underwent the specified interventions. Following the completion of experiment, the cardiomyocytes were treated with JC-1 (2 μM) according to the manufacturer’s protocol. Fluorescence imaging was examined using an IX-83 confocal microscope (Olympus, Japan).

### Lactic dehydrogenase activity

The H9C2 cardiomyocytes were seed in 6-well plates for 24 h. The cardiomyocytes were challenged with control siRNA or UCP2 siRNA for 48 h followed by H/R intervention. The NRG-1 postconditioning was executed immediately following the completion of hypoxic intervention. At 12 h after reoxygenation, the supernatant of cultured cardiomyocytes was collected to determine lactic dehydrogenase (LDH) activity using the commercial LDH activity assay kit (Elabscience Biotechnology Co., Ltd, Wuhan, China). The activity of LDH was determined following the manufacturer’s guidelines.

### Western blotting

At the end of experiment, both the I/R myocardium and H9C2 cardiomyocytes were collected for detecting the protein expression with western blotting. In short, both the I/R myocardium and H/R cardiomyocytes were lysed by the radio-immunoprecipitation assay buffer (Beyotime Biotechnology, Shanghai, China) with 1 mM phenylmethylsulfonyl fluoride and 1 mM protease inhibitor cocktail (MedChemExpress). The total protein concentration was determined with the bicinchoninic acid protein assay kit in accordance with the manufacturer’s guidelines (Beijing Solarbio Science & Technology Co., Ltd., China). The tissue homogenates and cell lysates were equally combined with 8-12% SDS-PAGE and transferred onto the polyvinylidene difluoride membranes (0.45 μm, Merck, Darmstadt, Germany). After washing with 0.05% Tween-20 in the Tris-buffered saline (TBST), the membranes were incubated at a room temperature condition for 2 h with a blocking solution containing 5% BSA and then incubated with specific primary antibody against IL-1β (Cell Signaling Technology, Danvers, Massachusetts, USA), LC3B (ABclonal Technology, Wuhan, China), UCP2 (Santa Cruz Biotechnology), PINK1 (Cell Signaling Technology) and α-tubulin (Santa Cruz Biotechnology) overnight at 4 °C. Following three times washed with TBST, the membranes were incubated with the HRP-conjugated secondary antibody (Santa Cruz Biotechnology) a room temperature condition for 1 h and subsequently visualized using the enhanced chemiluminescent HRP substrate (Millipore, Darmstadt, Germany). The signal was detected with the ChemiDoc XRS^+^ System (Bio-Rad, Hercules, California, USA) and quantified by the ImageJ Software (NIH).

### Statistical analysis

The normality of distribution for all parametric data was evaluated using the Kolmogorov–Smirnov test. Additionally, the Levene median test was utilized to evaluate the homogeneity of variance within the parametric data. All parametric data were presented as means ± standard deviations (SD) and the SPSS 26.0 (IBM, USA) was applied for statistical analyses. The unpaired Student’s *t*-test was used for between-group comparisons of data and one-way analysis of variance followed with the Bonferroni test was used for multi-group comparisons. *P*<0.05 was considered statistically significant.

## Results

### NRG-1 postconditioning alleviated myocardial I/R injury

As shown in *Fig. 1*, the I/R intervention resulted in significant infarct area (*Figure 1A and B*) and raised serum cTnI and CK-MB concentrations (*Figure 1C and D*), indicating the successful establishment of myocardial I/R injury model. Compared with the IRI group, administration of NRG-1 postconditioning prior to the reperfusion significantly reduced the infarct size and serum cTnI and CK-MB concentrations (*Figure 1A-D*). As compared with the Sham group, the IRI group presented the histological features of serious myocardial injury with HE staining, as evidence by irregular muscle bundles, massive ruptured muscle fibers and aggregation of neutrophils. However, these histological features were significantly mitigated in the NRG-1 group compared with the IRI group (*Figure 1E*). Similarly, the I/R intervention significantly upregulated IL-1β expression in the I/R myocardium, whereas NRG-1 postconditioning decreased IL-1β expression (*Figure 1F and G*). These results indicate that NRG-1 postconditioning produces a protection against myocardial I/R injury, with a significant inflammatory inhibition.

**Figure 1.**
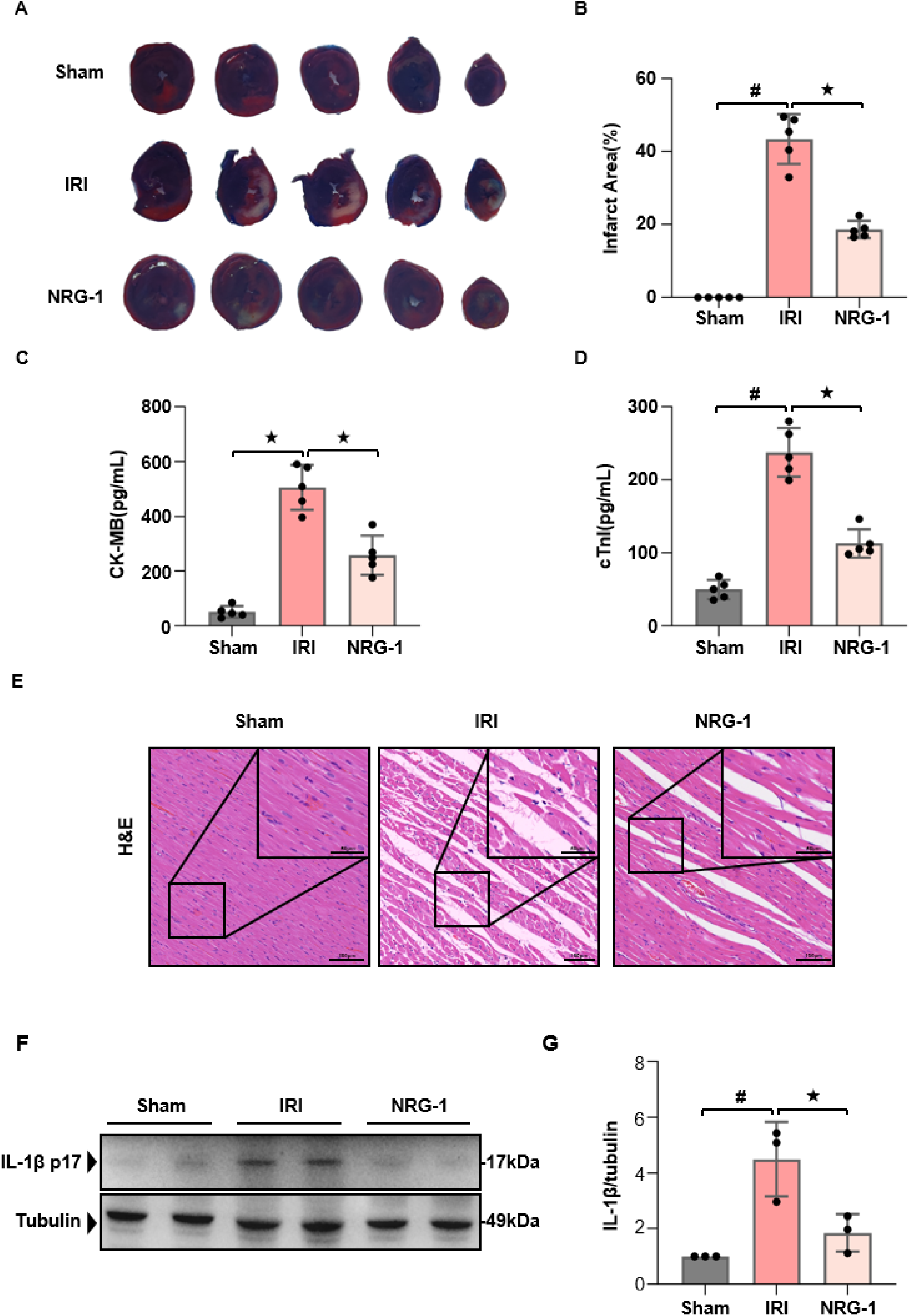
NRG-1 postconditioning attenuated myocardial I/R injury and inhibited myocardial inflammatory responses. (*A*) Representative appearances the I/R myocardium with Evans blue and TTC staining. (*B*) Quantitative analysis of infarct area (*n*=5). (*C, D*) Serum cTnI and CK-MB concentrations (*n*=5). (*E*) Representative pathological appearances of the I/R myocardium with H & E staining (magnification, ×20; inset, ×40; scale bar: 100/50 μm). (*F*) Representative western blots for interleukin-1 β (IL-1β, 17 kDa) and Tubulin (49 kDa, loading control) in the I/R myocardium. (*G*) Quantitative analysis of western blotting data for IL-1β in the I/R myocardium (*n*=3). The Unpaired Student’s *t*-test for between-group comparisons of data and one-way ANOVA followed with post hoc test (Bonferroni test) for multi-group comparisons. Data represent means ± SD. ^★^*P* < 0.05, ^#^*P* < 0.001

### The cardioprotection of NRG-1 postconditioning was associated with mitophagy activation

The experiment determined the differentially expressed genes and signaling pathways in the I/R myocardium of the IRI and NRG-1 groups using the RNA sequencing. As shown in *Figure 2*, the principal component analysis (PCA) displayed the similarities in the sample components of the IRI and NRG-1 groups, but the samples from different groups exhibited the distinct clustered distributions (*Figure 2A*). A total of 878 differentially expressed genes (DEGs) in the I/R myocardium were identified, comprising 455 up-regulated genes and 423 down-regulated genes (*Figure 2B*). Further KEGG pathway enrichment analysis of DEGs indicated that NRG-1 postconditioning significantly regulated the signaling pathways associated with mitophagy in the I/R myocardium (*Figure 2C*). Additionally, both Gene Set Enrichment Analysis (GSEA) and heatmap corroborated that NRG-1 postconditioning activated mitophagy in the I/R myocardium (*Figure 2D and E*). These data suggest that cardioprotection of NRG-1 postconditioning is associated with mitophagy activation in the I/R myocardium.

**Figure 2.**
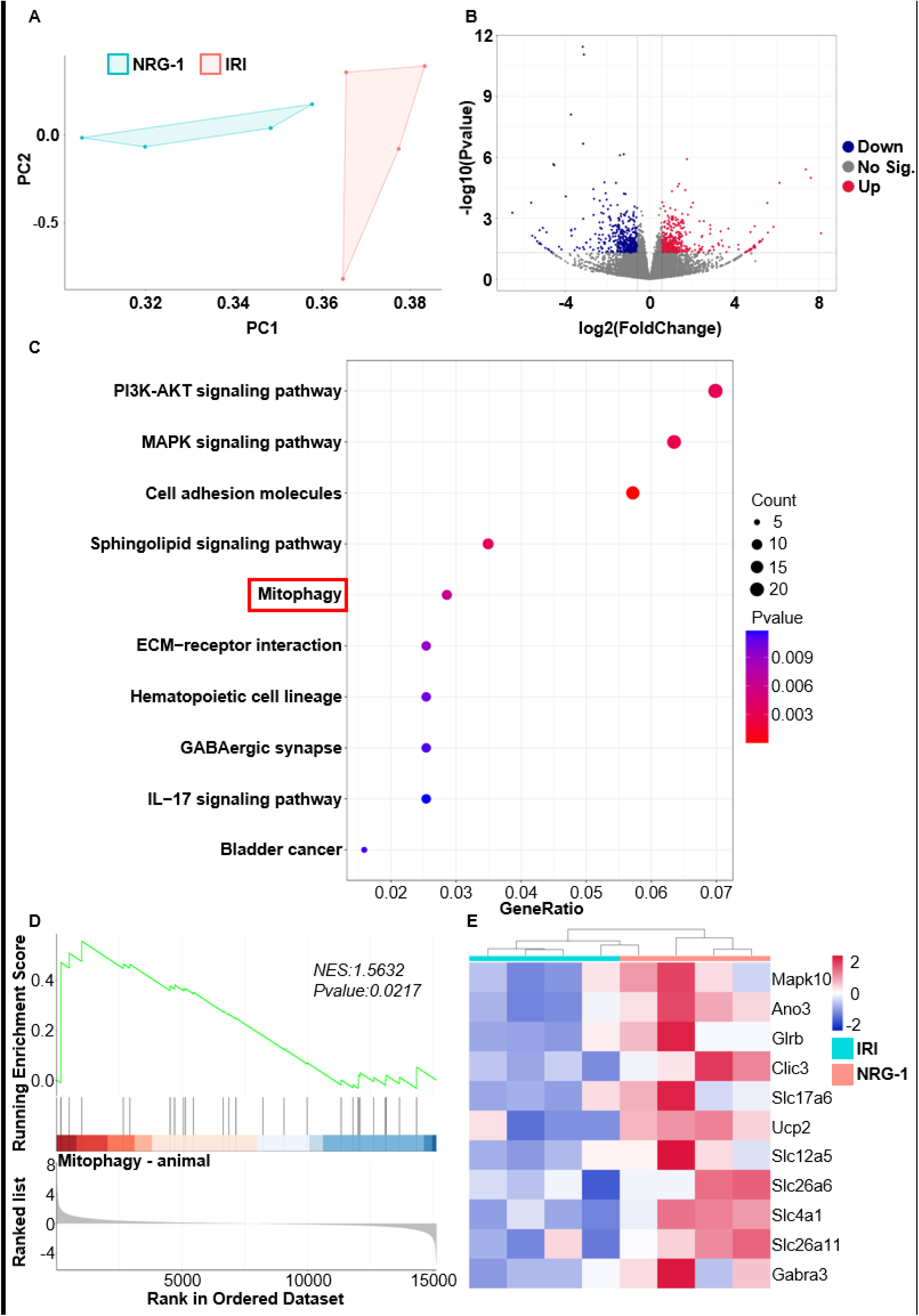
The cardioprotection of NRG-1 postconditioning was associated with mitophagy activation. (*A, B*) The PCA map of genes and volcano map of DEGs in the I/R myocardium from the IRI and NRG-1 groups (*n*=4, *P*<0.05). (*C*) The KEGG analysis of DEGs (*n*=4, *P*<0.05). (*D*) The GSEA analysis of mitophagy related genes with DEGs (*n*=4, *p* value is indicated on the figure). (*E*) The heat map of mitophagy related genes in the I/R myocardium from the IRI and NRG-1 groups (*n*=4).

### Inhibiting mitophagy eliminated cardioprotective effects of NRG-1 postconditioning

To elucidate the regulatory role of NRG-1 on mitophagy *in vivo*, mitophagy inhibitor Mdivi-1 was employed to assess its impact on the cardioprotective effects of NRG-1 postconditioning. In comparison to the Sham group, the IRI group appeared significant mitochondrial damages in the I/R myocardium, characterized by the pronounced mitochondrial swelling and disruption of intramitochondrial cristae with the TEM. Notably, NRG-1 postconditioning clearly attenuated the mitochondrial damages in the I/R myocardium, with damaged mitochondria being encapsulated by autophagic lysosomes, indicating the occurrence of mitophagy. However, Mdivi-1 treatment further exacerbated mitochondrial damages in the I/R myocardium and negated the mitophagy-activating effects of NRG-1 (*Figure 3A*).

**Figure 3.**
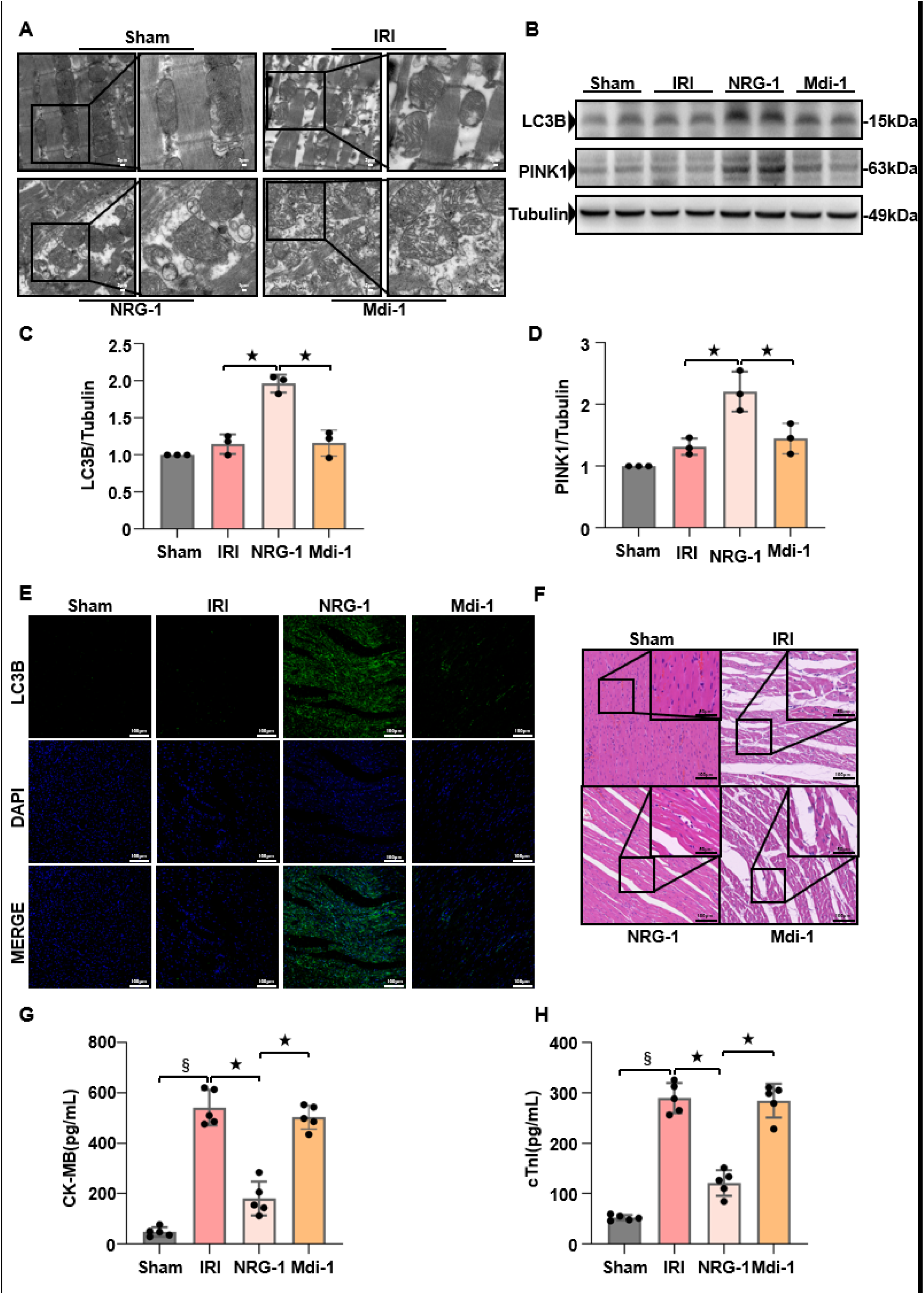
Inhibiting mitophagy eliminated the cardioprotective effects of NRG-1 postconditioning. (*A*) Representative TEM images of mitochondria in the I/R myocardium from different groups (magnification, ×5000; inset, ×10000; scale bar: 2/1 μm). (*B*) Representative western blots for LC3B (15 kDa), PINK1 (63 kDa) and Tubulin (49 kDa, loading control) in the I/R myocardium. (*C*, *D*) Quantification of western blotting data for LC3B and PINK1 in the I/R myocardium (*n*=3). (*E*) Representative confocal images of LC3B merged with DAPI in the myocardium (magnification, ×20; cale bar: 100 μm). (*F*) Representative myocardium pathological appearance of H&E staining (magnification, ×20; inset, ×40; scale bar: 100/50 μm). (*G*, *H*) Serum concentrations of cTnI and CK-MB (*n*=5). The Unpaired Student’s *t*-test for between-group comparisons of data and one-way ANOVA followed with post hoc test (Bonferroni test) for multi-group comparisons. Data represent means ± SD.^★^*P* < 0.05, ^§^*P* < 0.0001.

We further assessed the expressions of mitophagy-related proteins in the I/R myocardium. In comparison to the IRI group, NRG-1 postconditioning significantly enhanced the expression levels of LC3B and PINK1 in the I/R myocardium (*Figure 3B-D*), and increased the level of immunofluorescence-labeled LC3B in the I/R myocardium (*Figure 3E*). Nevertheless, Mdivi-1 treatment counteracted the enhancing effects of NRG-1 on these proteins associated with mitophagy (*Figure 3B-E*).

Compared with the IRI group, NRG-1 postconditioning significantly mitigated the myocardial I/R injury and lowered the serum cTnI and CK-MB concentrations. However, Mdivi-1 treatment eliminated the cardioprotective effects of NRG-1 postconditioning (*Figure 3F-H*). These results indicate that the cardioprotective benefits of NRG-1 postconditioning may be abolished by inhibiting mitophagy.

### The regulatory effect of NRG-1 on myocardial mitophagy was associated with UCP2

To determine the potential molecular mechanisms which NRG-1 regulates mitophagy in the I/R myocardium, we identified that 878 DEGs and 970 positively correlated genes (PCGs) were associated with mitophagy. The intersection of the two gene sets displayed 130 shared genes (*Figure 4A*). The correlation analysis between mitophagy and these genes indicated that the four genes exhibiting the strongest correlations were UCP2 (*R*=0.90, *P*<0.001), Kcnq5 (*R*=0.89, *P*<0.05), Camk2n2 (*R*=0.86, *P*<0.05), and Cdh1 (*R*=0.85, *P*<0.05), with UCP2 demonstrating the most robust correlation with mitophagy (*Figure 4B-E*). Then, the interactions between UCP2 and proteins associated with mitophagy were examined by constructing Protein-Protein Interaction Networks (PPI) using the STRING database. Among these proteins, UCP2 is known to interact with PINK1, MFN1/2 and VDAC1. Notably, UCP2 and PINK1 exhibited the strongest evidence for interaction, shown by an interaction score of 0.564 (*Figure 4F*). These data suggest that the regulatory effect of NRG-1 on myocardial mitophagy is associated with UCP2.

**Figure 4.**
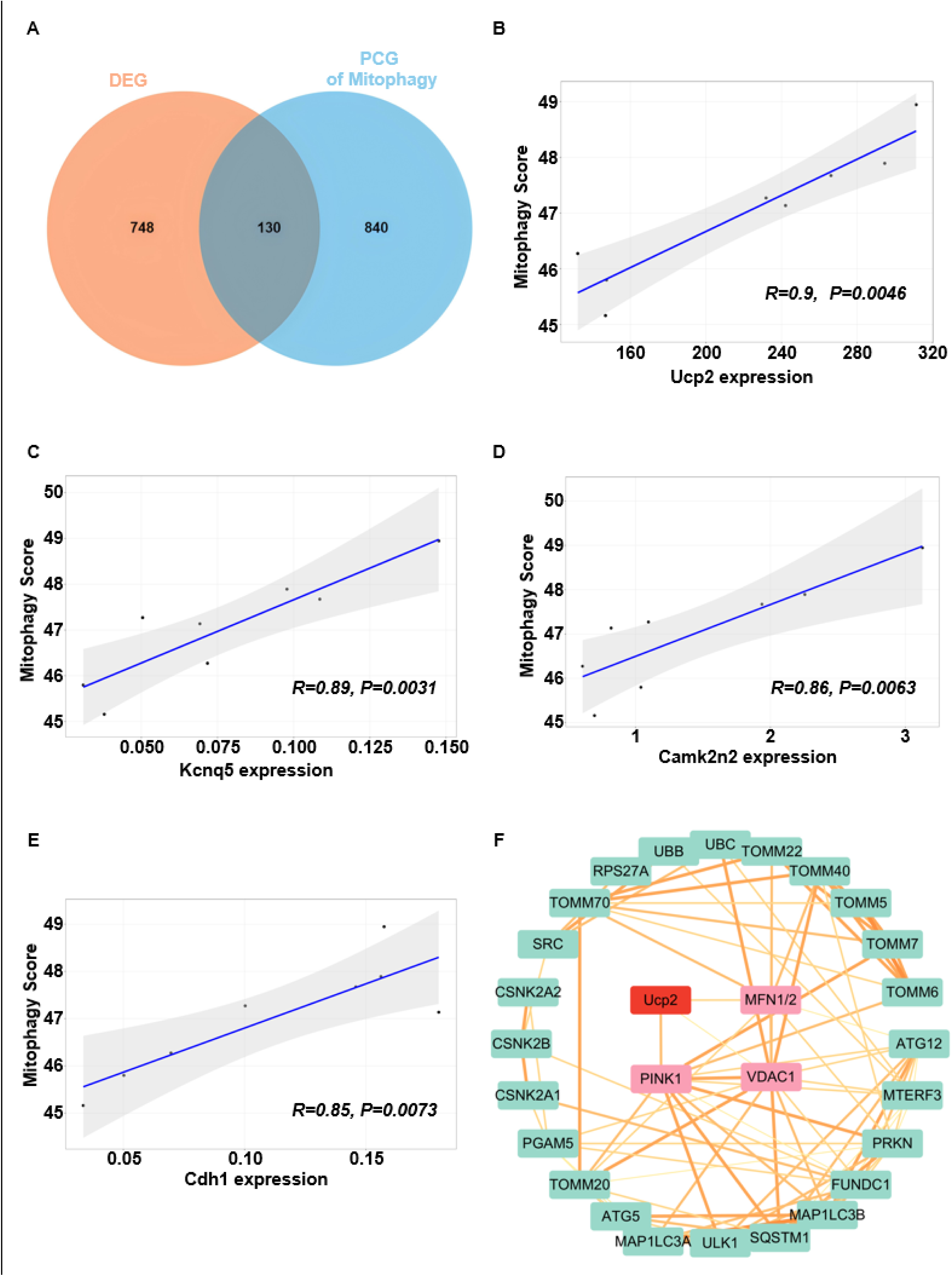
The regulatory effect of NRG-1 on mitophagy was associated with UCP2. (*A*) The Venn Diagram of the shared genes from the DEGs and PCGs associated with mitophagy (*n*=4). (*B*-*E*) The correlation analysis between shared genes and mitophagy (*n*=4, *p* value is indicated on the figure). (*F*)The PPI analysis between UCP2 and mitophagy-related proteins (the connection between the two indicates evidence of interaction, the thickness of the connection correlates with the strength of the evidence).

### The UCP2 was located at the upstream of myocardial mitophagy to mediate the regulatory role of NRG-1 postconditioning

As PPI analysis indicated a potential interaction between UCP2 and mitophagy, further *in vivo* experiment was carried out to validate the upstream/downstream relationships between UCP2 and mitophagy. As shown in *Figure 5*, compared with the I/R intervention, NRG-1 postconditioning significantly upregulated the UCP2 expression, and the expressions of mitophagy related protein LC3B and PINK1 in the I/R myocardium (*Figure 5A-D*). In comparison to NRG-1 postconditioning, Mdivi-1 treatment decreased the expression levels of LC3B and PINK1 in the I/R myocardium but did not change the UCP2 expression (*Figure 5A-D*). These results suggest that UCP2 is located at the upstream of mitophagy in the context of NRG-1-mediated activation of this process.

**Figure 5.**
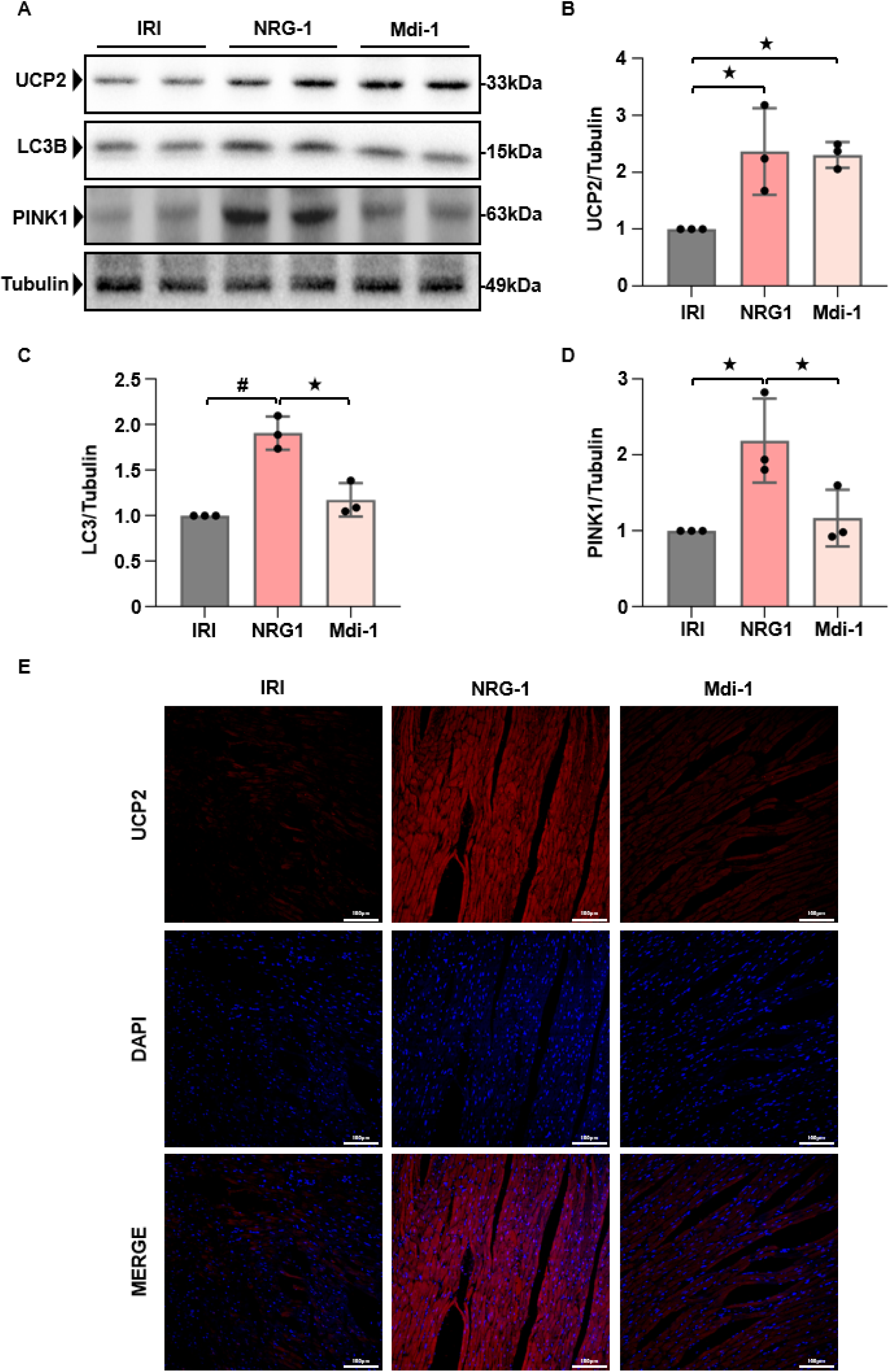
The UCP2 was located at the upstream of myocardial mitophagy to mediate the regulatory role of NRG-1 postconditioning. (*A*) Representative western blots for UCP2 (33 kDa), LC3B (15 kDa), PINK1 (63 kDa) and Tubulin (49 kDa, loading control) in the I/R myocardium. (*B*-*D*) The quantitative analyses of western blotting data for UCP2, LC3B and PINK1 in the I/R myocardium (*n*=3). (*E*) Representative confocal images of UCP2 merged with DAPI in the I/R myocardium (magnification, ×20; cale bar: 100 μm). The Unpaired Student’s *t*-test for between-group comparisons of data and one-way ANOVA followed with post hoc test (Bonferroni test) for multi-group comparisons. Data represent means ± SD.^★^*P* < 0.05, ^#^*P* < 0.001.

### NRG-1 postconditioning induced mitophagy in the cardiomyocytes subjected to H/R intervention

To further elucidate the potential molecular mechanism which NRG-1 postconditioning activated mitophagy in the I/R myocardium, an *in vitro* model of H9C2 cardiomyocytes receiving the H/R intervention was applied to observe the effects of NRG-1 on the UCP2 expression and mitophagy. As shown in *Figure 6*, compared with the H/R intervention, NRG-1 treatment significantly upregulated the expressions of UCP2 and PINK1 in the H/R cardiomyocytes (*Figure 6A-C*). The confocal microscopy revealed that NRG-1 treatment compared with the H/R intervention resulted in a significantly enhanced fluorescence intensity of JC-1 aggregates in the H/R cardiomyocytes (*Figure 6D*). These results preliminarily suggest that NRG-1 treatment activates mitophagy in the H/R cardiomyocytes, which is accompanied by an upregulated UCP2 expression.

**Figure 6.**
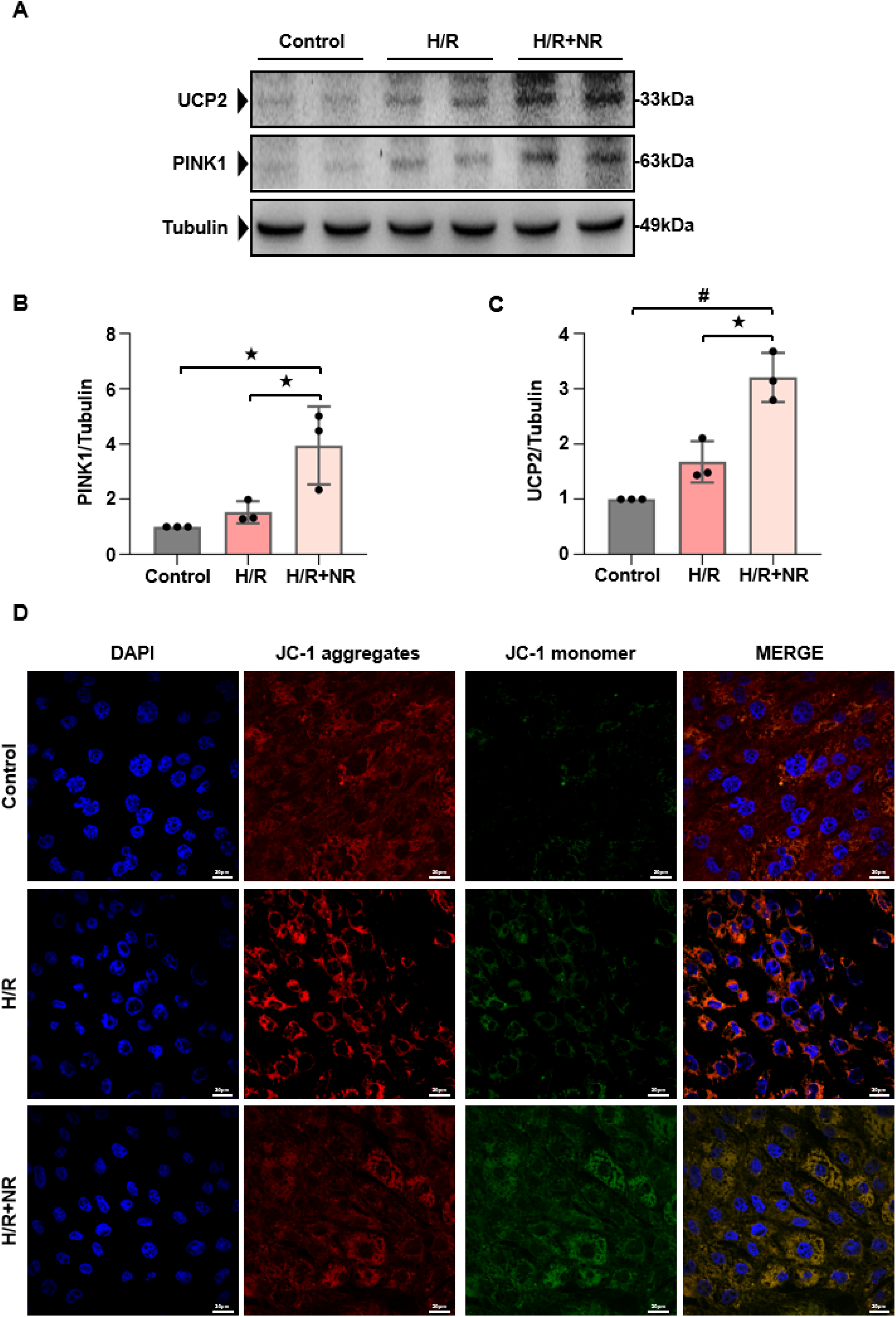
NRG-1 postconditioning induced mitophagy in the cardiomyocytes subjected to H/R intervention. (*A*) Representative western blots for UCP2 (33 kDa), PINK1 (63 kDa) and Tubulin (49 kDa, loading control) in the H/R cardiomyocytes. (*B*, *C*) The quantitative analyses of western blotting data for UCP2 and PINK1 in the H/R cardiomyocytes(*n*=3). (*D*) Representative confocal images of JC-1 staining merged with DAPI in the H/R cardiomyocytes (magnification, ×60; cale bar: 20 μm). The Unpaired Student’s *t*-test for between-group comparisons of data and one-way ANOVA followed with post hoc test (Bonferroni test) for multi-group comparisons. Data represent means ± SD. ^★^*P* < 0.05, ^#^*P* < 0.001.

### NRG-1 promoted the mitophagy in the H/R cardiomyocytes by upregulating the UCP2-PINK1-LC3B signaling pathway

To further verify the involvement of UCP2 in the activation of NRG-1 on mitophagy, the siRNA was applied to silence the UCP2 expression in the H/R cardiomyocytes and observe whether the mitophagy activation of NRG-1 could be abolished. As demonstrated in the *Figure 7A*, the UCP2 expression in the cardiomyocytes treated with NRG-1 was significantly reduced by the UCP2 siRNA, while no such reduction was observed with the negative control siRNA. The expression levels of mitophagy-associated proteins including PINK1 and LC3B were significantly elevated in the cardiomyocytes treated with NRG-1, but their changes were abolished by the treatment with UCP2 siRNA (*Figure 7A-D*). To assess the occurrence of mitophagy, the immunofluorescence staining was applied to label the LC3B and mitochondrial membranes, and the confocal microscopy was performed to observe the fluorescence colocalization. In the H/R cardiomyocytes treated with NRG-1, a significant fluorescence colocalization was observed, whereas this was diminished by the treatment with UCP2 siRNA (*Figure 7E*). The supernatant from cardiomyocytes subjected to the H/R intervention exhibited a significantly increased LDH activity, while the supernatant from cardiomyocytes treated with NRG-1 had a notably reduced LDH activity. Furthermore, the cardioprotective benefits of NRG-1 were decreased by the treatment with UCP2 siRNA (*Figure 7F*). Collectively, these data indicate that the signaling pathway involving the UCP2-PINK1-LC3B is crucial for the protective benefits of NRG-1 against cardiomyocyte H/R injury.

**Figure 7.**
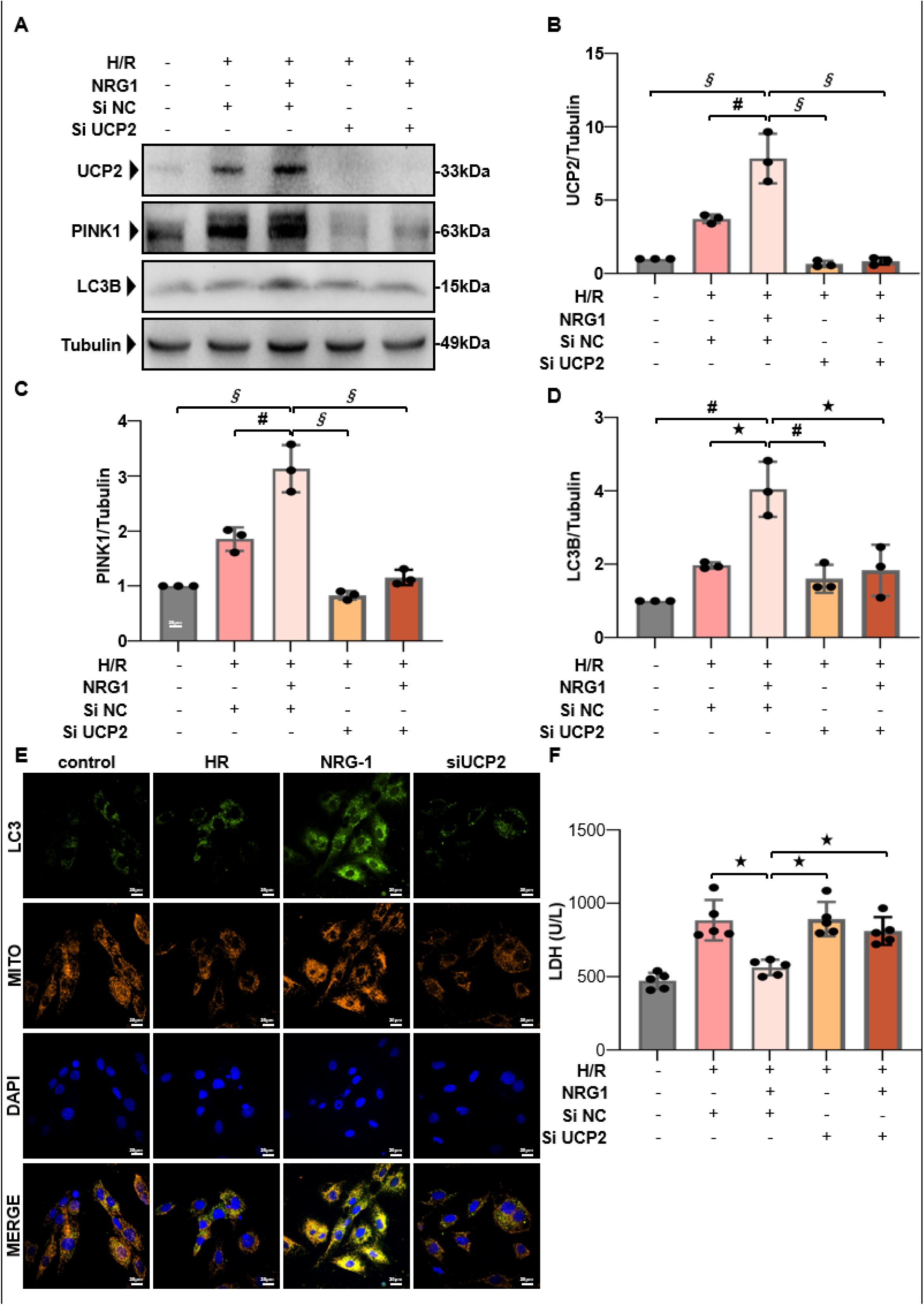
NRG-1 promoted mitophagy in the cardiomyocytes by upregulating the UCP2-PINK1-LC3B signaling pathway. (*A*) Representative western blots for UCP2 (33 kDa), PINK1 (63 kDa), LC3B (15 kDa) and Tubulin (49 kDa, loading control) in the H/R cardiomyocytes with or without UCP2 siRNA challenge. (*B*-*D*) The quantitative analyses of western blotting data for UCP2, PINK1 and LC3B in the H/R cardiomyocytes (*n*=3). (*E*) Representative confocal images of colocalization of LC3B and mitochondria and merged with DAPI in the H/R cardiomyocytes (yellow fluorescence indicates the occurrence of colocalization magnification, ×60; cale bar: 20 μm). (*F*) The LDH activity in the supernatants of cultured cardiomyocytes (*n*=5). The Unpaired Student’s *t*-test for between-group comparisons of data and one-way ANOVA followed with post hoc test (Bonferroni test) for multi-group comparisons. Data represent means ± SD. ^★^*P* < 0.05, ^#^*P* < 0.001, ^§^*P* < 0.0001.

### NRG-1 postconditioning attenuated myocardial I/R injury via the UCP2-PINK1-LC3B signaling pathway

In comparison to NRG-1 group, the treatment with UCP2 inhibitor genipin significantly decreased expression levels of UCP2, LC3B and PINK1 in the I/R myocardium (*Figure 8A-E*). Furthermore, the histological features of myocardial I/R injury were significantly mitigated in the NRG-1 group compared with the IRI and Genipin groups (*Figure 8F*). As compared with the NRG-1 group, the serum cTnI and CK-MB concentrations significantly reduced in the Genipin group (*Figure 8G and H*). These findings indicate that the protection of NRG-1 against myocardial I/R injury is achieved by activating mitophagy via the UCP2-PINK1-LC3B signaling pathway.

**Figure 8.**
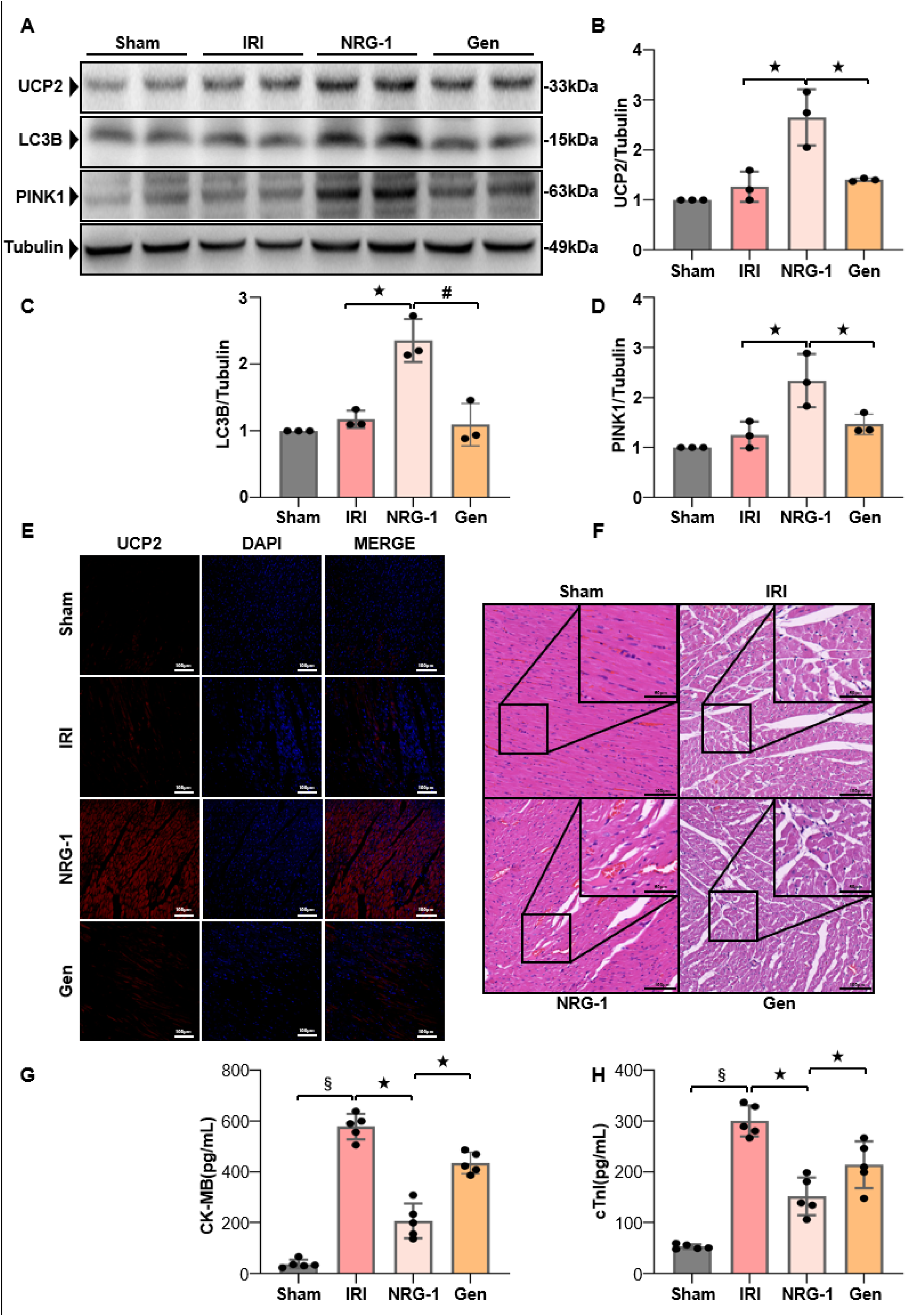
NRG-1 postconditioning attenuated myocardial I/R injury via the UCP2-PINK1-LC3B signaling pathway. (*A*) Representative western blots for UCP2 (33 kDa), LC3B (15 kDa), PINK1 (63 kDa) and Tubulin (49 kDa, loading control) in the I/R myocardium. (*B*-*D*) The quantitative analyses of western blotting data for UCP2, LC3B and PINK1 in the I/R myocardium (*n*=3). (*E*) Representative confocal images of UCP2 merged with DAPI in the I/R myocardium (magnification, ×20; cale bar: 100 μm). (*F*) Representative pathological appearances of the I/R myocardium with H & E staining (magnification, ×20; inset, ×40; scale bar: 100/50 μm). (*G*, *H*) Serum cTnI and CK-MB concentrations (*n*=5). The Unpaired Student’s *t*-test for between-group comparisons of data and one-way ANOVA followed with post hoc test (Bonferroni test) for multi-group comparisons. Data represent means ± SD. ^★^*P* < 0.05; ^#^*P* < 0.001; ^§^*P* < 0.0001.

## Discussion

NRG-1 is an endogenously produced mediator which is released from the endothelial cells during ischemia^26^. Available evidence indicates that NRG-1 can not only protect against myocardial I/R injury, but also it may serve as one of the key effectors of different cardioprotective interventions, such as the ischemia preconditioning^27^. Nevertheless, it is still unknown if mitophagy is involved in the cardioprotective benefits of NRG-1 and what the potential molecular mechanisms are involved. The main results of our experiment included: 1) NRG-1 postconditioning clearly attenuated myocardial I/R injury and upregulated the expression levels of LC3B and PINK1 in the I/R myocardium, while inhibiting myocardial mitophagy negated the cardioprotective benefits and decreased the effects of NRG-1 on these proteins; 2) Both the UCP2 and the PINK1 exhibited the strongest evidence for interaction through the PPI analysis, and the UCP2 siRNA abolished the up-regulating effects of NRG-1 on PINK1 and LC3B in the H/R cardiomyocytes. These findings suggest that the protection of NRG-1 against myocardial I/R injury is achieved by activating mitophagy via the UCP2-PINK1-LC3B signaling pathway.

The preconditioning strategies, such as ischemia preconditioning^28^, are the most effectively protective interventions for myocardial I/R injury, but their widespread application is limited due to the unpredictability of acute ischemic myocardial attack and the safety concerns in clinical practice^29^. The administration of pharmacological agents that share cardioprotective mechanisms with preconditioning may present a more viable alternative. It has been shown that NRG-1 postconditioning can provide a significant protection against myocardial I/R injury^9,18^. However, the detailed potential mechanisms remain unclear. In this experiment, thus, the RNA sequencing technique was employed to elucidate the specific mechanisms underlying the cardioprotective effects of NRG-1 postconditioning, and then mitophagy inhibitor was applied in the *in vivo* experiment and the UCP2 siRNA was administered in the *in vitro* experiment to validate the effects of NRG-1 on mitophagy and UCP2, respectively.

It has been shown that the expression level of IL-1β serves as a crucial indicator of myocardial inflammatory responses and cardiomyocyte damage during the I/R process^30^. In our experiment, NRG-1 significantly reduced the IL-1β expression in the I/R myocardium, suggesting that NRG-1 may inhibit the myocardial inflammatory responses during the I/R process. The previous work also demonstrates that NRG-1 postconditioning can result in anti-inflammatory effects by inhibiting pyroptosis^18^. In fact, mitochondrial dysfunction is a critical factor that triggers pyroptosis^31^. All of these findings hint that the anti-inflammatory effects of NRG-1 may be related to the modulation of mitochondrial function. However, there is the lack of experimental evidence supporting this hypothesis.

The MQC refers to the timely removal of damaged mitochondria and the reutilization of mitochondrial contents, which is very crucial for maintaining the normal function of cells^32,33^. The MQC consists of three main processes: mitochondrial biogenesis, dynamics (fusion and fission), and mitophagy. As a primary process of the MQC, mitophagy is responsible for the clearance of aging or dysfunctional mitochondria under physiological conditions, ensuring the maintenance of normal mitochondrial function^34^. In response to stress conditions, such as oxidative stress, nutrient deprivation, or pathogen infection, mitophagy is activated to eliminate damaged mitochondria, preventing cellular damage and death^35^. It has been shown that mitophagy plays a significant role in mitigating myocardial I/R injury^11,36^. Nevertheless, there are the limited researches that determine the direct influence of NRG-1 on the regulation of mitophagy in the context of myocardial I/R injury. By the RNA sequencing, this experiment showed there were the differentially expressed genes in the I/R myocardium between the IRI and NRG-1 groups. Furthermore, both the results of KEGG and GSEA analyses demonstrated that mitophagy was evidently enriched in the I/R myocardium following the NRG-1 treatment. Additionally, Mdivi-1 treatment counteracted the enhancing effect of NRG-1 on mitophagy in the I/R myocardium and reduced the cardioprotective benefits offered by NRG-1. Together with these findings, we deem that mitophagy activation is attributable to the cardioprotective effects of NRG-1 postconditioning.

The results of correlation analysis showed that among the DEGs, the UCP2 exhibited a strongest correlation with mitophagy. It is usually considered that the UCP2 can facilitate the immediate entry of protons into the mitochondrial matrix without engaging in ATP synthesis, thereby uncoupling oxidative phosphorylation, which leads to the reductions in the ATP synthesis and ΔΨm^37^. Nevertheless, a reduction in the ΔΨm is an important premise that triggers PINK1/LC3B-mediated mitophagy. Furthermore, it has been shown that overexpression of UCP2 may confer cardioprotective effects by inducing mitophagy, while the use of mdivi-1 treatment can negate the cardioprotective effects associated with UCP2 overexpression^38^. In this experiment, the PPI analysis between UCP2 and mitophagy showed that both UCP2 and PINK1 exhibited strongest evidence for interaction. The PINK1 is a key target that mediates the mitophagy. It has been demonstrated that the loss of PINK1 expression can result in mitochondrial dysfunction, thereby exacerbating the extent of myocardial damage associated with the I/R process^39^. In addition, there has been no study assessing the upstream and downstream regulatory relationship between the UCP2 and mitophagy. Our experiment demonstrated that the expression levels of UCP2 and mitophagy related proteins in the I/R myocardium were significantly upregulated by NRG-1 treatment. Especially, mitophagy inhibitor Mdivi-1 did not eliminate the upregulation of UCP2 in the I/R myocardium induced by NRG-1 treatment. Consequently, we consider that NRG-1 postconditioning may confer cardioprotection by regulating the UCP2/PINK1-mediated mitophagy.

The ΔΨm is a principal indicator for assessing mitochondrial function^40^and the ΔΨm alteration represent a critical component of the mechanisms underlying cardiomyocyte injury^41^. Furthermore, a reduction in the ΔΨm signifies the initiation of mitophagy^25^. By these variables, our *in vitro* experiment in the H/R cardiomyocytes further investigated the regulation of NRG-1 on the UCP2-mediated mitophagy. Our results showed that NRG-1 treatment markly enhanced the expression levels of UCP2 and PINK1 and decreased the ΔΨm in the H/R cardiomyocytes, indicating that NRG-1 treatment can activate the UCP2/PINK1-mediated mitophagy.

The impaired mitochondria can cause the decrease in the ΔΨm, which inhibits the translocation of PINK1 to the inner membrane of mitochondria. Subsequently, the PINK1 complex is bound to the phagocytic vesicle surface molecule LC3B, prompting the fusion of phagocytic vesicles with impaired mitochondria to form autophagosomes, thereby initiating mitophagy^42^. Our *in vitro* experiment in the H/R cardiomyocytes showed that NRG-1 treatment upregulated the expression levels of PINK1, UCP2 and LC3B, with the colocalization of LC3B and mitochondria in immunofluorescence staining. Nevertheless, these influences of NRG-1 were abolished by the intervention of UCP2 SiRNA. These results strongly support our hypothesis that NRG-1 can activate the mitophagy via the UCP2/PINK1/LC3B signaling pathway in the H/R cardiomyocytes.

To further verify our findings by the *in vivo* experiment, the treatment with UCP2 inhibitor genipin were also carried out in the rat model of myocardial I/R injury. The results demonstrated that genipin treatment not only obviated the expression upregulations of UCP2, PINK1 and LC3B in the I/R myocardium induced by NRG-1 postconditioning, but also significantly aggravated the myocardial I/R injury.

According to the findings of this experiment, we consider that the use of NRG-1 postconditioning to activate the UCP2/PINK1/LC3B-mediated mitophagy may be a useful strategy attenuating the myocardial I/R injury. Given that the function of MQC is to maintain a dynamic equilibrium within intracellular mitochondria, however, excessive activation of mitophagy can limit the intracellular energy supply and negatively impact cellular activity^43^, potentially exacerbating myocardial I/R injury^44^. Consequently, the moderate regulation of mitophagy is crucial for mitigating myocardial I/R injury. Due to the limitation of designing a single dose, however, this experiment cannot answer whether excessive NRG-1 postconditioning may lead to an imbalance in mitochondrial homeostasis. To achieve an optimal protection from NRG-1 postconditioning against myocardial I/R injury, further experiments are required to establish the appropriate dose and application time of this intervention.

In a word, our experiment demonstrates that NRG-1 postconditioning can produce a protection against myocardial I/R injury by enhancing the mitophagy through the UCP2/PINK1/LC3B signaling pathway. The NRG-1 postconditioning upregulates the expression level of UCP2 and decreases the ΔΨm, which activate the PINK1/LC3B-mediated mitophagy and then eliminate damaged mitochondria (*Figure 9*).

**Figure 9.**
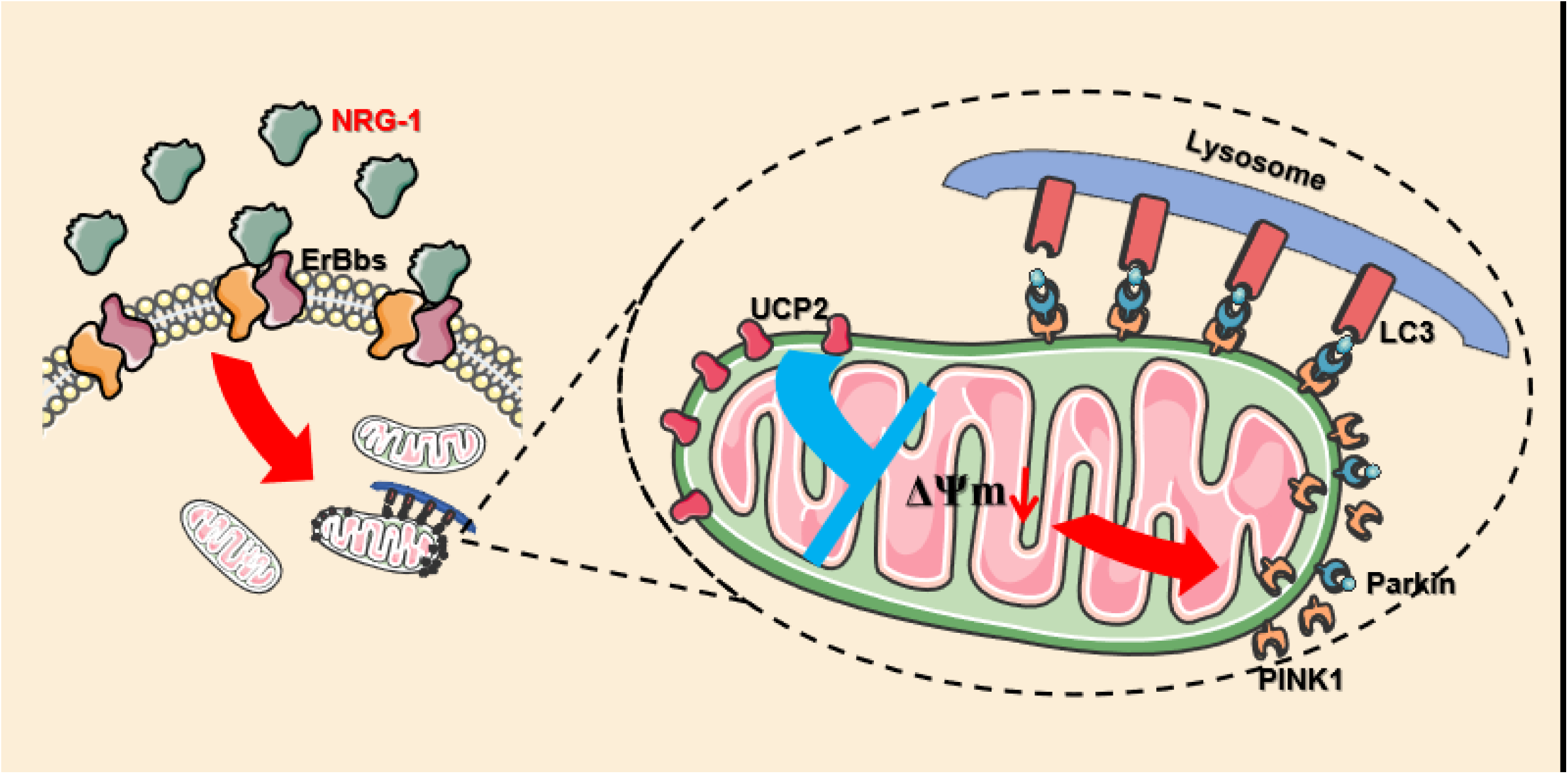
Schematic illustration exhibiting the molecular mechanism that NRG-1 activates mitophagy to provide a protection against myocardial I/R injury. The NRG-1 treatment upregulates the UCP2 expression in the mitochondria of cardiomyocytes. The UCP2 is able to reduce the mitochondrial membrane potential (MMP), which subsequently leads to the accumulation of the PINK1 complex on the outer membrane of mitochondria. This process induces the lysosomal surface recognizer LC3B to bind to the complex, ultimately resulting in the activation of mitophagy.

In conclusion, NRG-1 postconditioning can attenuate myocardial I/R injury by activating mitophagy through the UCP2/PINK1/LC3B signaling pathway. Our findings unveil a new insight into the potential mechanisms of myocardial I/R injury and provide a novel strategy for cardioprotective intervention.

## Acknowledgments

Sources of Funding

This study was supported by the National Natural Science Foundation of China (No. 81470019 to **F.S.X.**).

## Non-standard Abbreviations and Acronyms

AMI: acute myocardial infarction

I/R: ischemia/reperfusion

NRG-1: neuregulin-1

H/R: hypoxia/reoxygenation

MQC: mitochondrial quality control

MMP: mitochondrial membrane potential

LAD: left anterior descending

siRNA: small interfering RNA

AAR: ischemic area at risk

CK-MB: creatine kinase isoenzyme

cTnI: cardiac troponin I

LDH: lactic dehydrogenase

PCA: principal component analysis

DEGs: differentially expressed genes

TEM: transmission Electron Microscopy

GSEA: Gene Set Enrichment Analysis

PCGs: positively correlated genes

PPI: Protein-Protein Interaction Networks

## Novelty and Significance

### What is known?

- Endothelial cell-derived neuregulin-1 (NRG-1) may protecte against myocardial ischemia/reperfusion (I/R) injury and is involved in cardioprotective benefits of various interventions.
- Mitophagy is a crucial component of mitochondrial quality control (MQC), and its moderate activation can provide cardioprotective benefits.

### What new information does this article contribute?

- The cardioprotective effect of NRG-1 postconditioning is associated with the activation of mitophagy in the I/R myocardium.
- The activation of mitophagy by NRG-1 is associated with the upregulation of UCP2 expression.
- NRG-1 reduces mitochondrial membrane potential (MMP) by upregulating the expression of UCP2, which in turn triggers the PINK1/LC3B-mediated mitophagy.

Traditional treatments for myocardial I/R injury often face challenges in clinical practice. Therefore, it is more beneficial to identify pharmacological treatments that operate through similar mechanisms as these established methods. Mitophagy is a crucial aspect of MQC, and its moderate activation is essential for maintaining mitochondrial function. Our study demonstrates that NRG-1 can exert cardioprotective effects through the activation of mitophagy, thereby offering novel insights and a potential target for the treatment of myocardial I/R injury.

